# Large Neutral Amino acid uptake and mTOR activation within CD4+ T cells coordinate Type 2 immunity and host resistance to *Trichuris muris*

**DOI:** 10.1101/2020.11.30.405316

**Authors:** Maria Z Krauss, Kelly S Hayes, Ana Villegas-Mendez, Matthew R Hepworth, Linda V Sinclair, Kevin N Couper, Richard K Grencis

**Affiliations:** The Lydia Becker Institute of Immunology and Inflammation, Faculty of Biology, Medicine and Health, University of Manchester, Manchester, United Kingdom; Manchester Collaborative Centre for Inflammation Research, Division of Infection, Immunity and Respiratory Medicine, School of Biological Sciences, Manchester Academic Health Science Centre, University of Manchester, Manchester, UK; Wellcome Trust Centre for Cell-Matrix Research, University of Manchester, Manchester, United Kingdom; Cell Signalling and Immunology, University of Dundee, Dundee, UK

## Abstract

*Trichuris trichiura* (whipworm) is a gastrointestinal nematode that infects approximately 465 million people worldwide. *T. muris* is used as a tractable model for the human whipworm. In wild type mice, infection with a high dose of *T. muris* eggs leads to worm expulsion, which is dependent on a CD4+Th2 response and interleukin (IL-)13 production. It is known that T cells up-regulate glycolysis and uptake of substrates upon activation. The amino acid transporter SLC7A5 has been shown necessary for activation of mTORC1, a nutrient/energy/redox sensor critical for T cell differentiation into effector cells. We found that at the peak of the immune response to *T. muris*, mice lacking SLC7A5 in CD4+T cells have delayed worm expulsion, lower levels of IL-13, reduced pmTOR and glycolytic rates. However, at later stages of infection IL-13 levels partially recovered alongside resistance. The critical role of CD4+T cell metabolism *per se* and down-stream mTOR in CD4+T cells in resistance was shown in mice lacking mTOR in CD4+T cells, that failed to expel a high dose of parasites and developed chronic infection. Our study shows that mTOR is essential for effective functioning of T cells during whipworm infection and that deletion of Slc7a5 significantly delays worm clearance.

## INTRODUCTION

Over the last few years it has been established that cellular metabolism dictates the function of immune cells (1). Remodelling metabolism is especially important for T cells. By trafficking through the body, they are exposed to different sites and nutrient availability and when activated they need to rapidly proliferate and secrete cytokines (2, 3). The highly increased metabolic demand of activated T cells needs to be supported by increased uptake of nutrients such as lipids, glucose and amino acids (4). mTOR is a master regulator of T cell metabolism that integrates activating cues received including via the TCR, co-stimulatory signals and cytokines, with signals from the environment (2, 5). mTOR then coordinates the activation of downstream signalling pathways that control nutrient uptake, glycolysis, mitochondrial biogenesis and fatty acid oxidation (2). Upon activation of naïve CD4+ T cells, mTOR promotes upregulation of glycolysis (6). mTOR^-^ CD4 T cells were shown to undergo normal activation but failed to differentiate into Th1, Th2 and Th17 subsets (7). mTOR signalling occurs via two distinct complexes, mTORC1 and mTORC2 (8). The main function of mTORC1 is to control cellular growth and proliferation and the complex is required for Th1 and Th17 differentiation (9, 10). mTORC2 regulates cellular functions such as actin reorganization and is essential for Th2 cell differentiation (5, 9). Although both mTORC1 and mTORC2 play a major role in regulating cellular metabolism, little is known about the importance of T cell mTOR in type 2 infectious models (6, 10).

Slc7a5 (also known as large neutral amino acid transporter 1; LAT1) is the main large neutral amino acid transporter in T cells and was described as essential for their subset differentiation and expansion (4). Slc7a5 is an antiporter that can transport several amino acids, with an important role to transport leucine and methionine, essential neutral amino acids (11). Intracellular leucine is required for mTOR activation by coordinating mTOR positioning on lysosomal membranes and RAG GTPase activity (12). Methionine deprivation was also shown to partially ablate mTOR activity in T cells (13). Previous work has also shown the importance of Slc7a5 for Th1 and Th17 responses (4) although its role in Th2 controlled immunity is unknown.

Th2 responses have co-evolved in order to mediate protective immune responses to large extracellular organisms, such as gastrointestinal helminths. Soil-transmitted helminths (STH), including *Ascaris lumbricoides* (roundworm), *Trichuris trichiura* (whipworm), *Necator americanus* and *Ancylostoma duodenale* (hookworms) infect over 1 billion people worldwide. Infections are concentrated in sub-Saharan Africa, Asia, and Latin America, in areas where sanitation is poor and malnourishment is frequent and are recognised as Neglected Tropical Diseases (14). Infections are acquired early in life, chronic infection is the norm through to adulthood with parasites placing significant metabolic demands on the host (15, 16). *Trichuris muris* is a natural parasite of mice and is extensively utilised as a laboratory model for the study of human whipworm infection, *T*. *trichiura* (17) with remarkable similarity between species (18) and capacity to model both resistance and susceptibility. Resistance to *T. muris* is readily generated under experimental conditions and is dependent on a CD4+ Th2 response and production of interleukin (IL-)13, again in common with other murine models of STH, such as *Nippostrongylus brasiliensis* and *Heligmosoides polygyrus* (19, 20).

Here, we show that during acute *T. muris* infection the metabolic profile of CD4+ T cells is altered to promote worm expulsion. Generation of protective immunity is associated with elevation of expression of nutrient transporters and increased glycolytic rates in the CD4+T cells population. Using conditional knock out mice (mTOR^fl/fl^CD4-Cre mice), we show that T cell mTOR is essential for resistance against *T. muris*. Furthermore, deletion of Slc7a5 in CD4+T cells reduces mTOR activity and significantly delays the generation of protective immunity and worm expulsion. Taken together the data supports the hypothesis that CD4+Th2 cells are sensitive in terms of their metabolic demands (in this case amino acids) during intestinal helminth infection and changes in their capacity to respond appropriately can significantly alter their ability to clear pathogens. This has implications not only for other intestinal helminth infections but also for the wider field of Th2 mediated immunological disease at mucosal surfaces.

## RESULTS

When infected with a high dose of *T. muris* eggs, WT C57BL/6 mice develop a Th2 response, which expels the parasites through increasing epithelial cell turnover, muscle contractility and mucus production (21–23). Effective resistance to *T. muris* is dependent upon the generation of a strong and rapid CD4 T cell IL-13 response, to expel the worm burden before immunomodulation by the parasite and chronic infection becomes established (24).

### CD4 T cells change their metabolism upon *T. muris* infection

To assess the metabolic changes in CD4 T cells upon *T. muris* infection, C57BL/6 mice were infected with a high dose of infective eggs. CD71, the transferrin receptor was previously shown to be upregulated in activated T cells (25, 26), as well as CD98, which supports T cell proliferation by complexing with large amino acid transporters (27). Three weeks post infection, the peak time of cytokine production in this system, CD4 T cells from the mesenteric lymph node (MLN) showed increased expression of the metabolism related markers CD98 and CD71 and the proliferation marker Ki67 (Fig. 1a). Extracellular flux analysis of CD4+T cells isolated from infected mice showed up-regulated glycolytic metabolism in comparison to CD4+T cells from uninfected mice (Fig. 1b). CD4 T cells up-regulated both glycolytic rates and glycolytic capacity upon infection (Fig. 1c).

**Figure 1.**
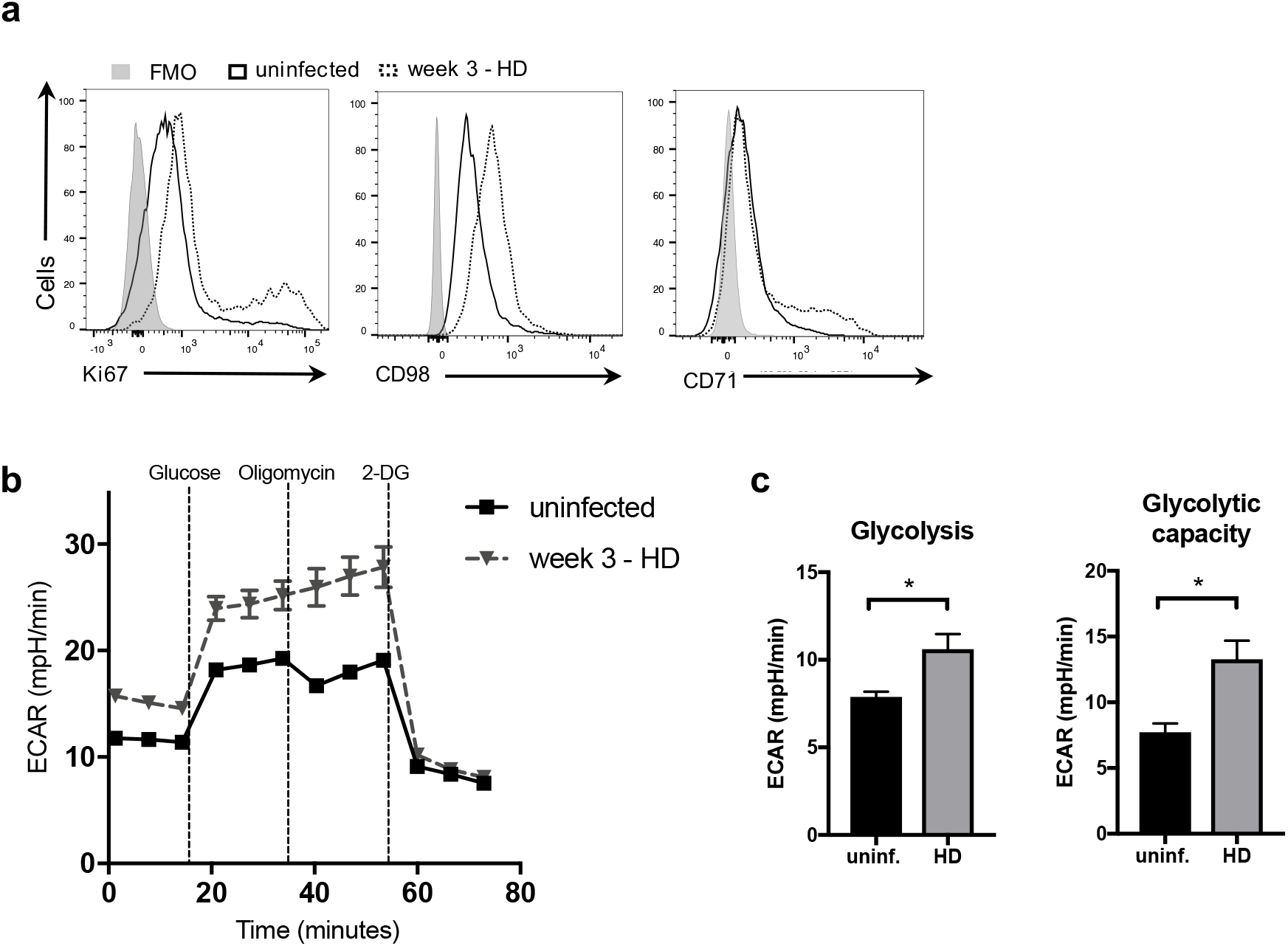
*T. muris* infection alters expression of nutrient trans porters and cell metabolism in CD4+ T cells. (a) Flow cytometric analysis of Ki67, CD98 and CD71 in CD4+ T cells from the mesenteric lymph node after 21 days of high dose *T. muris* infection in C57BL/6 mice. (b) Extracellular flux analysis of CD4+ T cells from MLN of high dose infected and naïve C57BL/6 mice. (c) Glycolytic rate and glycolytic capacity from CD4+ T cells from MLN of naïve and week three infected C57BL/6. Data are representative of two independent experiments with 5 mice per group.

### T cell mTOR is required for resistance against *T. muris*

mTOR is a master regulator of T cell metabolism and in order to define its importance in host protection, mTOR^fl/fl^CD4-Cre mice and mTOR^fl/fl^ control mice were infected with a high dose *T. muris* infection. mTOR^fl/fl^CD4-Cre mice failed to expel a high dose *T. muris* infection, even after 5 weeks (Fig. 2a). mTOR^-^ CD4+ T cells from MLN taken 3 weeks after infection (p.i) showed lower expression of CD44, CD71 and CD98 when compared to WT CD4+ T cells at this time point (Fig. 2c). However, overall CD4+ T cell counts in MLN from mTOR^fl/fl^CD4-Cre mice and mTOR^fl/fl^ mice were comparable (Fig. 2b). MLN cells from infected mTOR^fl/fl^CD4-Cre mice failed to produce IL-13 after re-stimulation with parasite E/S antigen (Fig. 2d). Parasite-specific IgG1 is detected in serum of wild type (WT) mice in response to *T. muris* infection (28). Serum levels of parasite-specific IgG1 found in infected mTOR^fl/fl^CD4-Cre mice were significantly reduced as compared to the levels found in infected mTOR^fl/fl^ mice (Fig. 2e). Levels of parasite-specific IgM, however, were found at similar levels in sera from infected mTOR^fl/fl^CD4-Cre mice and mTOR^fl/fl^ mice, indicating a potential failure of T cells to support B cell class switching or to produce the appropriate class switching cues (Fig. S1). To investigate further the impairment of IgG1 observed when mTOR was deleted in T cells, we also assessed the follicular helper T cell population upon infection in mTOR^fl/fl^CD4-Cre and mTOR^fl/fl^ mice. The PD-1+ CXCR5+ Tfh population was absent in infected mTOR^fl/fl^CD4-Cre mice (Fig. 2f). Expression of Bcl-6 was also found reduced in CD4+ T cells from these mice (Fig. S2). This confirms previous reports showing that mTOR is required for antibody class switching and Tfh differentiation (29, 30). We next assessed the impact of the deletion of mTOR on the metabolism of CD4+ T cells during infection, using extracellular flux analysis. mTOR^-^ CD4+ T cells not only failed to up-regulate glycolysis upon infection (Fig. 3a-b), but also showed lower levels of mitochondrial respiration when compared to WT CD4+ T cells (Fig. 3c-e). The metabolic phenotype of mTOR^-^ CD4+ T cells from infected mice was more quiescent-like than WT CD4+ T cells from infected mice (Fig. 3c). Despite having very high worm burdens for long periods of time, mTOR^fl/fl^CD4-Cre mice did not present with any aberrant pathologies (Fig.S3).

**Figure 2.**
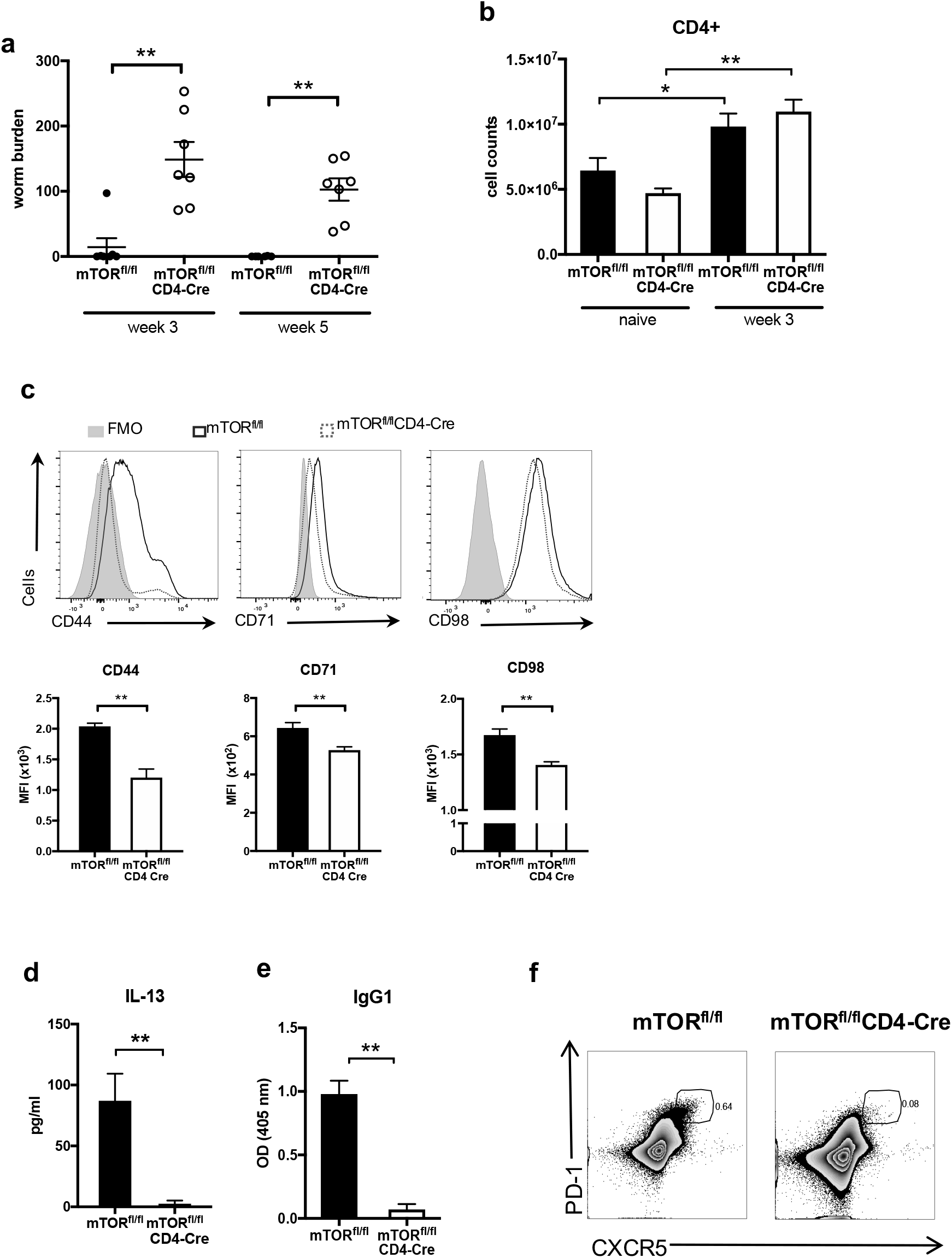
T cell mTOR is required for immunity to *T. muris*. (a) Worm burden from mTOR^fl/fl^ and mTOR^fl/fl^CD4-Cre mice after 3 and 5 weeks of high dose *T. muris* infection. (b) Number of CD4+ T cells in the mesenteric lymph nodes of naïve and high dose infected mTOR^fl/fl^ and mTOR^fl/fl^CD4-Cre mice, analysed by flow cytometry. (c) Flow cytometry analysis of CD44, CD71, CD98 and Ki67 in MLN cells from infected mTOR^fl/fl^ and mTOR^fl/fl^CD4-Cre mice. (d) MLN cells from mTOR^fl/fl^ and mTOR^fl/fl^CD4-Cre mice infected for 3 weeks were re-stimulated with parasite secretory/excretory product for 32 hours. Secretion of IL-13 was measured by cytometric bead array. (e) Levels of parasite-specific IgG1 in sera from week 3 infected mTOR^fl/fl^ and mTOR^fl/fl^CD4-Cre mice. (e) (f) Flow cytometric analysis of follicular helper T cells (PD-1+ CXCR5+) in infected mTOR^fl/fl^ and mTOR^fl/fl^CD4-Cre mice. (a) Graph shows individual replicates of animals for two independent experiments combined (n=3-4). (b-f) Data are representative of two independent experiments with 4-5 mice per group. * indicates p value ≤0.05, ** indicates p value≤ 0.01 (unpaired t test).

**Figure 3.**
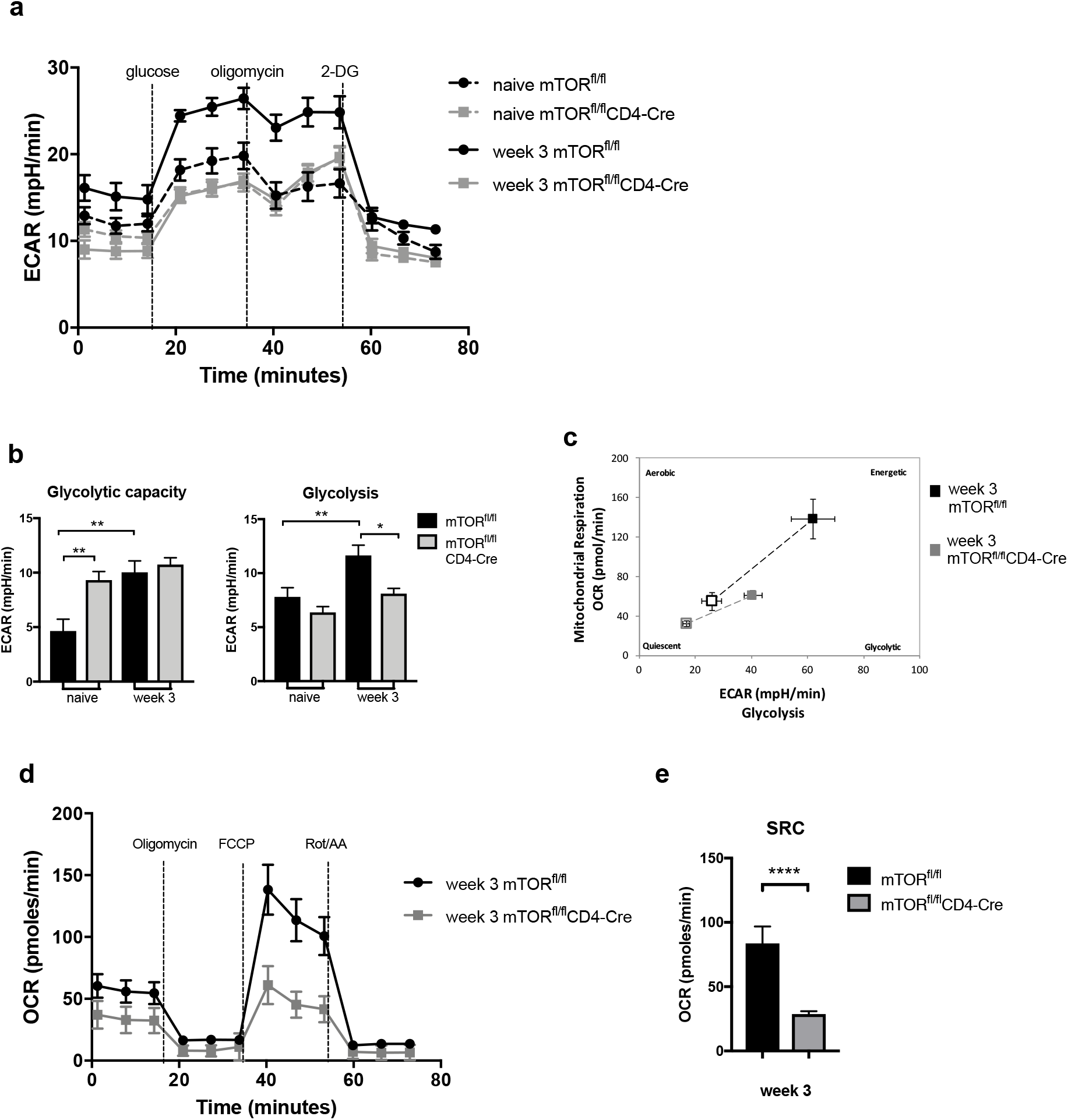
T cells fail to upregulate the glycolytic pathway upon *T. muris* infection in the absence of mTOR. (a) CD4+ T cells were isolated from MLNs of naïve and week 3 infected mTOR^fl/fl^ and mTOR^fl/fl^CD4-Cre mice using magnetic labelling. CD4+ T cells were subjected to a glycolytic stress test. (b) Quantification of glycolytic capacity and glycolysis levels of CD4+ T cells from mTOR^fl/fl^ and mTOR^fl/fl^CD4-Cre mice obtained with extracellular flux analysis. (c) Cell energy phenotype from CD4+ T cells isolated from MLN of infected mTOR^fl/fl^ and mTOR^fl/fl^CD4-Cre mice. Empty squares represent basal metabolic levels, filled squares represent stressed measurements (after injection of oligomycin and FFCP). (d) Mitochondrial stress test from CD4+ T cells from MLNs of week 3 infected mice. (e) Spare respiratory capacity of CD4+ T cells from mTOR^fl/fl^ and mTOR^fl/fl^CD4-Cre mice obtained with extracellular flux analysis. Cells from each mouse were individually isolated and all groups were submitted to the extracellular flux analysis simultaneously to allow comparisons. Graphs show mean ±SEM and data are representative of two independent experiments with 3-4 mice per group. * indicates p value ≤0.05, ** indicates p value≤ 0.01 (Kruskall-Wallis followed by Dunn’s multiple comparison test).

### Deletion of Slc7a5 in T cells leads to impairment of the Th2 response 3 weeks post infection

We questioned the nutrient transport systems that were required for mTOR activation and CD4+ T cell metabolic reprogramming during *T. muris* infection. Previous studies have shown that *Slc7a5* is required for T cell differentiation and clonal expansion, e.g. for transporting leucine that is required inside the cell for mTORC activation and for sustaining c-Myc expression (4, 31, 32). To assess the uptake of large neutral amino acids by CD4+T cells following *T. muris* infection kynurenine uptake was assessed *ex vivo*. Kynurenine uptake is an accurate proxy to measure LNAA uptake, and uptake has been shown to be SLC7A5 dependent in murine T cells (33). The data shows that following infection, activated CD4+T cells in both MLN and cecum utilise SLC7A5 to transport amino acids, and the addition of excess leucine competitively blocks the uptake of kynurenine to comparable levels seen using the system L blocker BCH (Fig.4a). To assess the importance of SLC7A5 for a type 2 response, Slc7a5^fl/fl^CD4-Cre mice (4) and Slc7a5^fl/fl^ control mice received a high dose infection of *T. muris*. (34, 35). We assessed infection of Slc7a5^fl/fl^CD4-Cre mice and in their littermate controls after 3 and 5 weeks to measure the peak cytokine responses and progression of the infection to patency respectively. Slc7a5^fl/fl^CD4-Cre mice showed significantly higher worm burdens than Slc7a5^fl/fl^ control mice 3 weeks after a high dose infection of *T. muris* (Fig. 4b). Slc7a5^fl/fl^CD4-Cre mice also showed reduced numbers of CD4+ T cells in the MLN (Fig. 4c). Frequencies of the activation marker CD44 in CD4+ T cells from MLNs of Slc7a5^fl/fl^CD4-Cre mice were slightly higher than those found in Slc7a5^fl/fl^ mice. However, percentages of CD71 and CD98 and the proliferation marker Ki67 in these CD44hi CD4+ T cells were found to be lower in the absence of T cell Slc7a5 at 3 weeks p.i. (Fig. 4d). Re-stimulation of MLN cells from infected mice with parasite secreted/excreted (E/S) antigen revealed that production of IL-13 was abrogated in Slc7a5^fl/fl^CD4-Cre mice (Fig. 4e). Serum levels of parasite-specific IgG1 were much lower in infected Slc7a5^fl/fl^CD4-Cre mice when compared to levels found in infected Slc7a5^fl/fl^ control mice (Fig. 4f). CD4+ T cells from Slc7a5^fl/fl^CD4-Cre mice showed lower phosphorylation of mTOR (S2448) in CD4+ T cells from MLNs (Fig. 4g).

**Figure 4.**
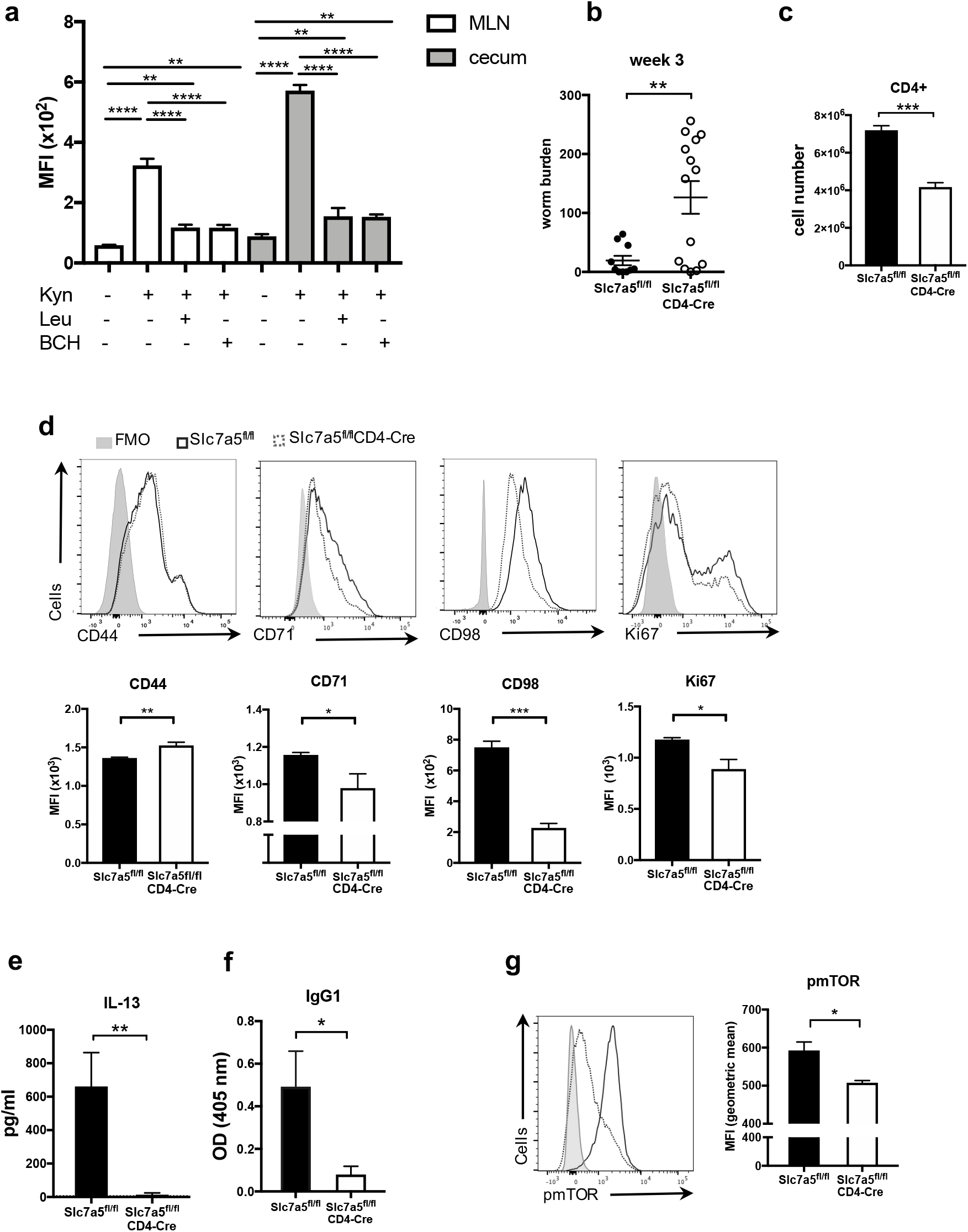
CD4+ T cells transport kynurenine via the system L transporter SLC7A5 during *T. muris* infection and Slc7a5^fl/fl^CD4-Cre mice exhibit impaired Th2 response three weeks after high dose *Trichuris muris* infection compared with wild-type (Slc7a5^fl/fl^) mice. (a) Kynurenine (200μM) uptake in CD4+ T cells from week 3 high dose infected C57BL/6 mice in the presence or absence of leucine (5mM) and BCH (10mM). (b) Number of *T. muris* larvae found in the caecum and proximal colon of Slc7a5^fl/fl^CD4-Cre and wild-type mice three weeks after high dose infection. (c) CD4 expression in cells from MLN was assessed by flow cytometry. (d) Flow cytometric analysis of CD44 in CD4+ T cells and of CD71, CD98 and Ki67 in CD44hi CD4+ T cells in MLN from infected Slc7a5^fl/fl^ and Slc7a5^fl/fl^CD4-Cre mice. (e) MLN cells from infected Slc7a5^fl/fl^ and Slc7a5^fl/fl^CD4-Cre mice were re-stimulated with parasite excretory/secretory product for 32 hours and IL-13 levels in cell supernatant were assessed by cytometric bead array. (f) Serum levels of parasite-specific IgG1 in Slc7a5^fl/fl^ and Slc7a5^fl/fl^CD4-Cre mice three weeks after *T. muris* infection. (g) Flow cytometic analysis of phosphorylated mTOR in CD4+ T cells from infected Slc7a5^fl/fl^ and Slc7a5^fl/fl^CD4-Cre mice. Graphs show mean ±SEM and data are representative of 2-3 independent experiments with 3-5 mice per group. * indicates p value ≤0.05, ** indicates p value≤ 0.01, ***indicates p value≤ 0.001, **** indicates p value< 0.0001. (a) paired t-test, (b-g) unpaired t-test.

### Worm expulsion and IL-13 production are partially rescued in Slc7a5^fl/fl^CD4-Cre mice at later stages of infection

When assessed after 5 weeks of a high dose *T. muris* infection, Slc7a5^fl/fl^CD4-Cre mice showed similar worm burdens to Slc7a5^fl/fl^ control mice, with most of the animals having expelled infection (Fig. 5a). At this time point, MLNs from Slc7a5^fl/fl^CD4-Cre mice had comparable numbers of CD4+ T cells to those from Slc7a5^fl/fl^ mice (Fig. 5b). Frequencies of CD44, CD71 and Ki67 CD4+ T cells were also found to be similar in MLN from infected Slc7a5^fl/fl^CD4-Cre and infected Slc7a5^fl/fl^ mice. Expression of CD98, however, remained much lower in CD4+ T cells from Slc7a5^fl/fl^CD4-Cre after 5 weeks of infection (Fig. 5c). Production of IL-13 by MLN cells upon re-stimulation with parasite E/S antigen was found to be similar between Slc7a5^fl/fl^CD4-Cre and Slc7a5^fl/fl^ mice after 5 weeks of infection (Fig. 5d). Also, serum levels of parasite-specific IgG1 were no longer different between both groups at this time point (Fig. 5e). Data from this later time point demonstrated that deletion of Slc7a5 in T cells significantly delays worm expulsion and the type 2 response but does not induce complete susceptibility to a high dose infection of *T. muris*.

**Figure 5.**
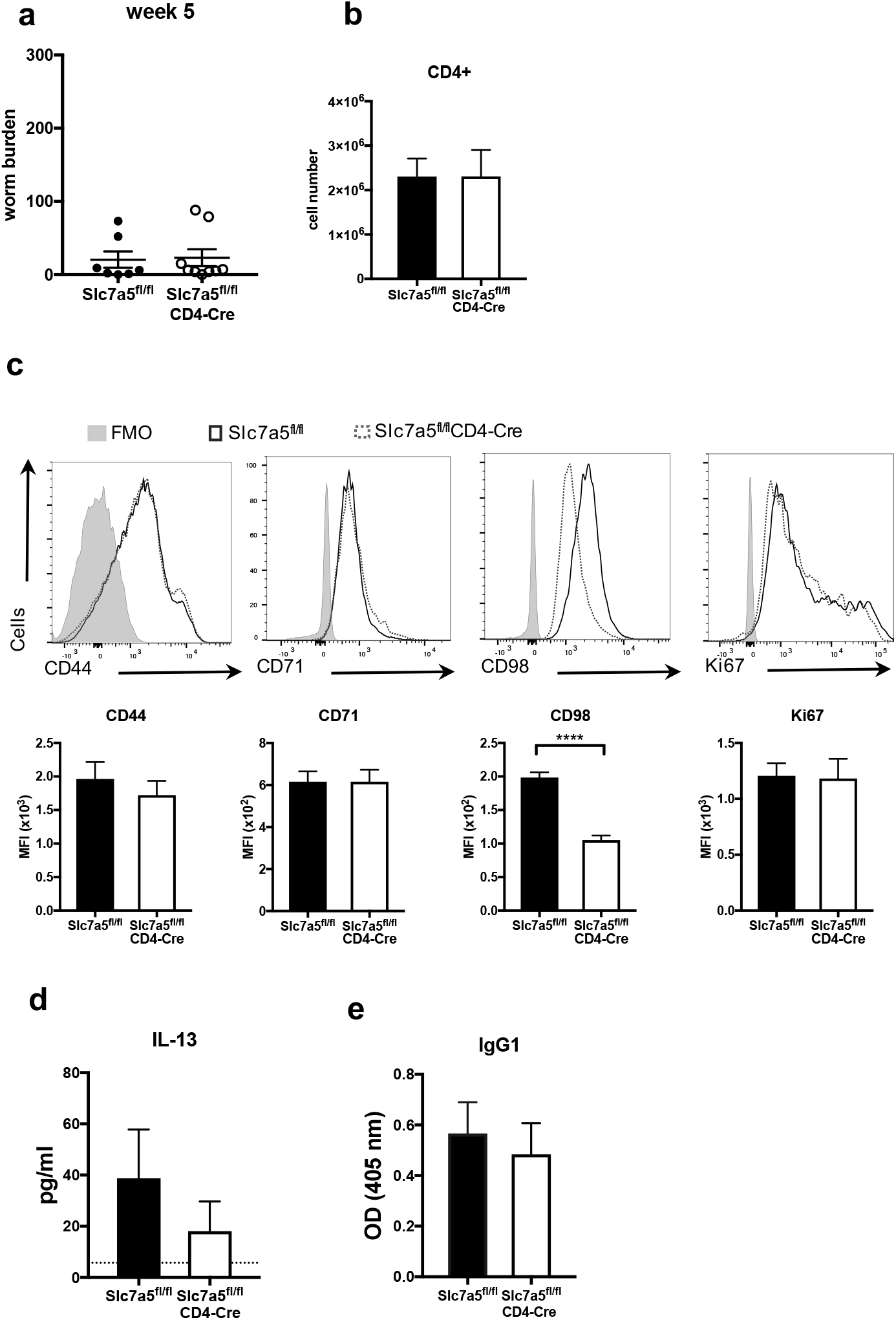
CD4+ T cells overcome the lack of Slc7a5 at the later stage of *T. muris* infection. (a) Worm burden after 5 weeks of high dose *T. muris* infection in Slc7a5^fl/fl^ and Slc7a5^fl/fl^CD4-Cre mice. (b) Numbers of CD4+ T cells in MLN after 5 weeks of high dose infection. (c) Flow cytometry analysis of CD44 in CD4+ T cells and of CD71, CD98 and Ki67 in CD44hi CD4+ T cells in MLN CD4+ T cells from 5-week-infected Slc7a5^fl/fl^ and Slc7a5^fl/fl^CD4-Cre mice. (d) MLN cells from infected Slc7a5^fl/fl^ and Slc7a5^fl/fl^CD4-Cre mice were incubated with the excretory/secretory antigen from the parasite for 32 hours. Levels of secreted IL-13 in cell supernatants were measured by cytokine bead array. Dotted line represents IL-13 secretion by cells from uninfected mice. (e) Serum levels of parasitespecific IgG1 from 5-week-infected Slc7a5^fl/fl^ and Slc7a5^fl/fl^CD4-Cre mice. (a) Graph shows individual replicates of animals for 2 independent experiments combined (n=4-5). (b-e) Graphs show mean ±SEM and data are representative of two independent experiments with 4-5 mice per group. **** indicates p value< 0.0001 (unpaired t-test).

### CD4+ T cells that lack Slc7a5 show lower glycolytic rates at the earlier stage of *T. muris* infection

Previous studies have shown that Slc7a5 was required for mTOR activation and expression of glucose transporters and, consequently, to support glycolysis (11, 31). Using extracellular flux analysis we investigated if deletion of Slc7a5 in CD4+ T cells from MLN resulted in impaired metabolic shift during infection. CD4+ T cells were isolated from infected Slc7a5^fl/fl^CD4-Cre and infected Slc7a5^fl/fl^ mice and subjected to a glycolytic stress test. At week 3 p.i., CD4+ T cells from littermate control mice showed higher glycolysis and glycolytic capacity when compared to those from Slc7a5^fl/fl^CD4-Cre mice (Fig. 6a). Levels of glycolytic capacity were also lower in CD4+ T cells from naïve Slc7a5^fl/fl^CD4-Cre mice when compared to those from naïve littermate control mice. After 5 weeks of *T. muris* high dose infection, Slc7a5^-^ CD4+ T cells showed similar glycolytic capacity and higher glycolytic levels to littermate control mice CD4+ T cells (Fig 6b). We therefore confirmed the metabolic impairment of the CD4+ T cells at the early stages of infection.

**Figure 6.**
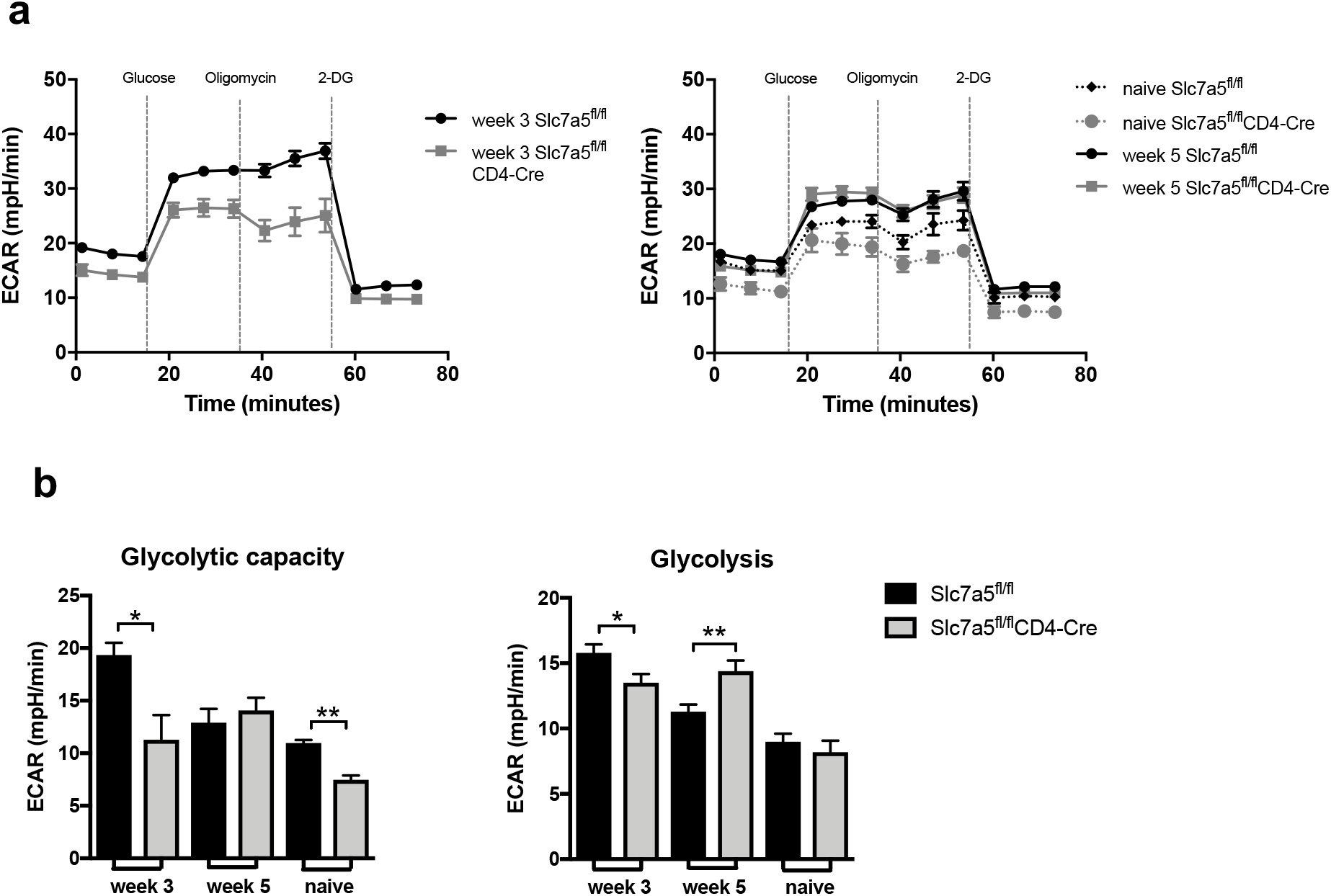
CD4+ T cells lacking Slc7a5 show impaired glucose metabolism 3 weeks post *T. muris* infection. (a) CD4+ T cells from infected and naïve Slc7a5^fl/fl^ and Slc7a5^fl/fl^CD4-Cre mice were isolated using magnetic beads and subjected to a glycolytic stress test. (b) Shows glycolytic capacity and glycolytic levels of CD4+ T cells from Slc7a5^fl/fl^ and Slc7a5^fl/fl^CD4-Cre mice obtained with extracellular flux analysis. Cells from each mouse were individually isolated and all groups were submitted to the extracellular flux analysis simultaneously to allow comparisons across time points. Graphs show mean ±SEM and data are representative of two independent experiments with 3-4 mice per group. * indicates p value ≤0.05, ** indicates p value≤ 0.01 (Mann-Whitney test).

### CD4+ T cells from mice chronically infected with *T. muris* show depressed metabolism

Our data show that a depression of glycolytic rates in CD4+ T cells from mice normally capable of expelling *T. muris* is associated with a reduction in host protective capabilities suggesting that CD4+ T cell metabolism plays a key role in the capacity of these cells to carry out their effector roles efficiently. As GI nematode infections are normally found as long-lived chronic infections i.e. unable to clear the parasites, it was of interest, therefore to examine the metabolism of CD4+ T cells taken from chronically infected mice. To investigate this a low dose infection was given to C57BL/6 mice, that progresses to patency. Previous work has shown that susceptibility to *T. muris* is associated with the generation of a CD4+Th1 response, not a Th2 response (28). Figure 7 (a-b) shows that CD4+T cells from infected mice show a significantly elevated glycolytic capacity at day 21 post infection consistent with this time point being the peak of Th1 and IFN-γ activity, but by day 35 post infection the glycolytic capacity was markedly depressed. Moreover, CD4+ T cell mitochondrial respiration was also depressed during chronic infection (Fig. 7 c-d) suggesting metabolic activity more akin to naïve CD4+ T cells, despite ongoing exposure to parasite antigen.

**Figure 7.**
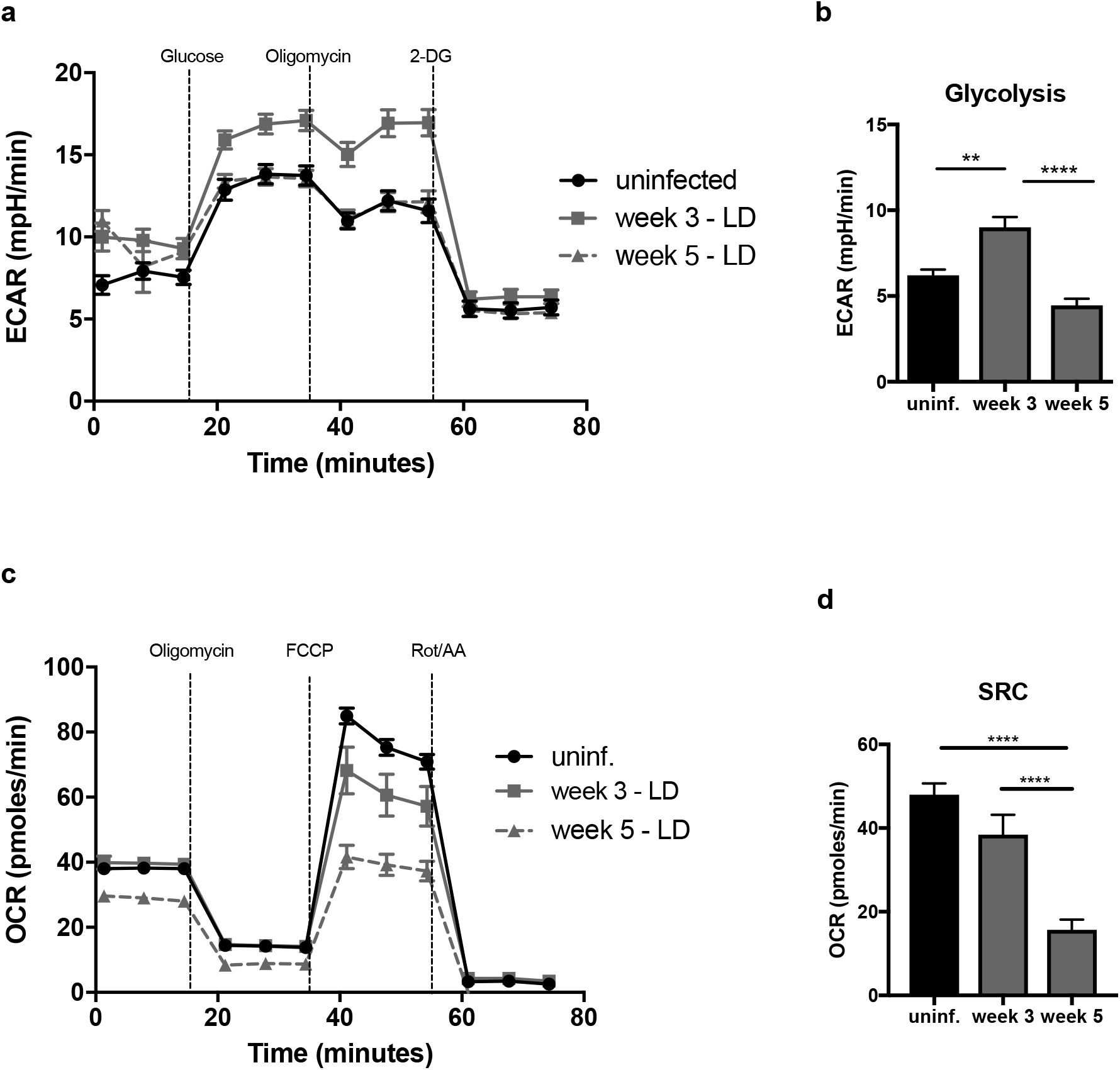
Chronic *T. muris* infection alters CD4+ T cell metabolism. (a) Extracellular flux analysis of CD4+ T cells from the mesenteric lymph nodes after 21 and 35 days of low dose infection in C57BL/6 mice, glycolytic stress test. (b) Glycolytic levels from figure 7a. (c) Mitochondrial stress test from CD4+ T cells from mesenteric lymph nodes at day 21 and 35 post low dose *T. muris* infection. (d) Spare respiratory capacity relative to figure 7c. Cells from each mouse were individually isolated. Graphs show mean ±SEM and data are representative of two independent experiments with 5 mice per group. ** indicates p value≤ 0.01, **** indicates p value< 0.0001 (Kruskall-Wallis followed by Dunn’s multiple comparison test).

## DISCUSSION

In this study we demonstrated that deletion of the amino acid transporter Slc7a5 delays expulsion of a high dose *T. muris* infection and that T cell mTOR is required for resistance. Assessment of the immune response at an early stage of infection revealed impairment of T cell function upon deletion of Slc7a5 in T cells, which included abrogation of IL-13 and IgG production. Slc7a5 can transport the large neutral amino acid leucine and intracellular leucine was previously shown to be essential for mTOR activation (36, 37). It was then hypothesised that the abrogation of T cell function observed in infected Slc7a5^fl/fl^CD4-Cre mice was due to a lack of intracellular leucine required for mTOR activation. Phosphorylation of mTOR was found decreased in Slc7a5 deficient CD4+ T cells at the early stage of infection suggesting that a lack of intracellular amino acids required for mTOR activation may be responsible. Also, at the earlier stages of infection CD4+ T cells lacking Slc7a5 exhibited a muted up-regulation of glycolysis compared to wild type CD4+ T cells. Slc7a5^fl/fl^CD4-Cre mice showed significantly reduced production of parasite-specific IgG during the earlier stages of infection, similarly to that observed in mTOR^fl/fl^CD4-Cre mice suggesting that Slc7a5 is important for T cell helper function. At later time points of infection, however, Slc7a5^fl/fl^CD4-Cre mice eventually generated an immune response which, although restricted, was sufficient to promote worm clearance. The possibility that Slc7a5^fl/fl^CD4-Cre mice started to re-express Slc7a5 was ruled out by RT-PCR *Slc7a5* analysis in CD4+ T cells from the later stages of infection (Fig. S4). This suggests compensatory large amino acid transporter mechanisms may be operating. CD98 can form heterodimers not only with Slc7a5 (LAT1), but also with Slc7a6 (y+LAT2), Slc7a7 (y+LAT1) and Slc7a8 (LAT2) (27). In this case, given that expression of CD98 remains low during the later phase, it is likely that the compensatory mechanism is at least partially independent of CD98 and of these LAT systems. It is possible that there are multiple amino acid transporting systems importing large neutral amino acids at the later stages of infection or that cells are using another amino acid instead of leucine. It appears that upon infection, T cells that lack Slc7a5 take some time to upregulate other transporter(s) that are capable of compensating for the absence of Slc7a5. After the compensatory transporters are in place, the cell is functional and Slc7a5^fl/fl^CD4-Cre mice can expel *T. muris* as fast as Slc7a5^fl/fl^ mice when re-challenged suggesting that once an effective Th2 response has been primed the absence of Slc7a5 is not critical (Fig. S5). Thus, early nutrient sensing by T cells together with availability of amino acids and glucose are key to the generation of efficient responses to *T. muris*. It is noteworthy that mice fed a low protein diet showed impaired resistance to a high dose *T. muris* infection (38). Intestinal nematode infection has multiple effects upon the metabolic phenotype of the host tissue (16) and nematodes themselves require the same resources for development and rapid growth. Indeed, transcriptomic analysis of *T. muris* confirms expression of parasite amino acid transporter genes including a large amino acid transporter and parasite glucose transporter genes in the larval stages that are present during the first 21 days of infection (18). Moreover, the bacillary band, a modified region of the cuticle found in *T. muris* and other parasitic nematodes that occupy an intestinal epithelial niche has also been shown to readily absorb glucose from the environment (39). Taken together, it is reasonable to suggest the parasites themselves are competing with host tissue including T cells for these valuable resources during establishment of infection.

Gastrointestinal helminth infections are regarded as chronic infections, with individuals harbouring infections for extended periods of time. Low dose *T. muris* infections allow us to assess the metabolism of CD4+T cells during susceptibility. The data clearly show that at day 21 post infection, the peak time of the generation of Th1 IFN-g response in the MLN (34), that CD4+ T cells show elevated glycolytic activity/capacity reflecting their activation. Clearly, low larval numbers do not impose a significant drain on available nutrients at this time. Adult worms, however, are particularly metabolically demanding and by day 35 post infection, CD4+T cells within the MLN show significantly depressed metabolic activity assessed by both glycolysis and mitochondrial respiration with profiles more similar to naïve cells despite being exposed to large quantities of parasite derived antigens. We may speculate that this is of benefit to the parasite/host relationship with the diminished metabolic capacity of the CD4+ T cells reflecting a poorly supportive environment for T cell activation reducing the pathological potential of Th1 cells in the intestine. It is noteworthy that T regulatory cell numbers are also depressed during low dose chronic *T. muris* infection (40, 41).

We demonstrate that mTOR plays a major role in T cell mediated immunity against *T. muris*. The deletion of mTOR in T cells had a severe impact on the immune response to *T. muris* and made mice completely susceptible to high dose infections. mTOR^fl/fl^CD4-Cre mice showed abrogated production of IL-13, impaired Tfh differentiation and antibody class switching and absence of CD4+ T cell accumulation in the large intestine (Fig. S6). As shown by data from the extracellular flux analysis, mTOR appears to not only be required for upregulation of aerobic glycolysis in T cells, but also for increased mitochondrial respiration during *T. muris* infection. This is in line with previous studies using other cell types that showed that mTOR can also promote mitochondrial activity (42, 43). The role of mTOR during intestinal infection has received attention with T cell mTOR essential for resistance to *Citrobacter rodentium* infection in mice (44), although in this case, for the generation of efficient host protective Th1 responses in the intestinal mucosa. Studies of the protozoan parasite, *Tritrichomonas muris* added a futher level of complexity showing that intestinal epithelial cell specific mTORC1 was important in the initiation of Type 2 immune responses (45).

Increased glycolytic levels were observed in CD4+ T cells upon infection and support a role for glycolysis in immunity against *T. muris*. This is in line with several previous studies which have described aerobic glycolysis a requirement for cytokine production in T cells (46, 47). Tibbit *et al* recently showed that Th2 cells are heavily reliant on glycolysis for their differentiation and function and that blocking glycolysis *in vivo* using 2-DG impairs the production of IL-13 by Th2 cells (48). Our work confirms the correlation between glycolysis and IL-13 production and reinforces that upregulation of glycolysis is dependent on mTOR. Overall, data presented here have demonstrated that CD4+ T cell mTOR acts as major regulator of the immune response to *T. muris*. Deletion of mTOR in T cells results in failure to upregulate metabolic pathways, which leads to major impairment of T cell activation and function and therefore susceptibility to *T. muris*. Moreover, deletion of Slc7a5 in T cells leads to considerable impairment of the effector function T cells in the early stage of infection. This reinforces that amino acid transport and availability sensing are a key step in licensing mTOR activation. T cells lacking Slc7a5, however, partially recover function later on infection and promote parasite expulsion. The finding that alteration of a single amino acid transporter could significantly influence the generation of protective immunity to an intestinal helminth confirms the importance of immune cell metabolism following infection. The present work demonstrates the fragile equilibrium of nutrient availability and nutrient transporters that exists for efficient T cell effector function at mucosal sites and it is intriguing to speculate that individuals with mutations in amino acid transporters that affect efficient transport may be compromised in their capacity to generate rapid CD4 T cell IL-13 responses. In endemic areas of gastrointestinal nematode infection, infected individuals are often malnourished. It is reasonable to speculate, for example, that low protein diets could contribute to making hosts more susceptible to helminths and potentially to other kinds of infection, due to compromised metabolic activity of mucosal immune cells.

## METHODS

### Animals

Slc7a5^fl/fl^CD4-Cre mice were obtained from Prof. Doreen Cantrell (University of Dundee, UK) (4). mTOR^fl/fl^CD4-Cre mice were originally purchased from JAX (USA). Animals were maintained in individually ventilated cages with 65% humidity at 21-23°C. Mice were used at 6-10 weeks of age and infected and naïve mice were cohoused. Mice were sex and age matched for experiments. Mice were kept in accordance with the regulations of the Home Office Scientific Procedures Act (1986) and were euthanized by CO_2_ inhalation.

### Infections and worm counts

Parasite maintenance and extraction of the E/S antigen were performed as described previously (49). For high dose infections, mice were infected with approximately 300 infective *T. muris* eggs in ddH2O via oral gavage. For low dose infections, mice were given 20 infective eggs in ddH_2_O via oral gavage. Worm burdens were assessed at 21 or 32-35 days post infection. For worm burden determination, the caecum and proximal colon were removed and cut longitudinally in a Petri dish with water. The intestinal contents were removed, the epithelium was scraped using curved forceps and the worms were individually removed and counted under a dissecting microscope.

### Cell culture and cytokine analysis

MLN cells were collected and re-stimulated for 32 hours as previously described (50). Secreted concentrations of IL-13 and IFN- γ were measured from cell supernatant using a cytometric bead array (CBA, BD Biosciences, UK) according to the manufacturer’s instructions.

### Parasite specific antibody analysis

Serum levels of parasite-specific antibodies were measured by ELISA as described previously (49).

### FACS

Flow cytometry experiments were performed on a BD LSRII or Fortessa, and analysed using FlowJo software. The following antibodies were used for flow cytometry: anti-CD4 (GK1.5, Biolegend), anti-CD98 (RL388, Biolegend), anti-PD-1 (29F.1A12, Biolegend), anti-CD71 (R17217, Biolegend), anti-Ki67 (B56, eBioscience), CXCR5 (L138D7, Biolegend), anti-pmTOR (MRRBY, eBioscience), anti-CD44 (IM7, Biolegend). Intracellular staining was performed using Fixing/Permeabilization Foxp3 kit (eBioscience). For assessment of phosphorylated mTOR, cells were fixed with 4% paraformaldehyde and permeabilized with methanol. Kynurenine uptake assays were performed as described previously (33).

### Extracellular flux analysis (Seahorse)

CD4+ T cells were isolated from MLNs using magnetic positive selection (L3T4 beads, Miltenyi) according to the manufacturer’s instructions. Cellular metabolism was assessed using a Seahorse Bioscience XF96e Extracellular Flux Analyzer. Approximately 200000 isolated CD4+ T cells were adhered to the Seahorse plate using Cell-Tak (Corning) and cultured in Seahorse media for 30min – 1h. ECAR was measured in XF media (modified DMEM containing 2 mM L-glutamine) under basal conditions, in response to 10mM glucose, 2μM oligomycin and 100mM 2-DG. OCAR was measured under basal conditions, after injection of 2μM oligomycin, 1.5μM FCCP and 0.5μM rotenone + 0.5μM antimycin A (all Sigma). If necessary, data was normalized by cell number after analysis.

## ACKNOWLEDGMENT

We thank members of the R.K.G. and K.N.C. laboratories (University of Manchester) for scientific discussions and some experimental assistance. We thank Doreen Cantrell (University of Dundee) for the Slc7a5fl/flCD4-cre mice and the Flow Cytometry Facility (University of Manchester) for the support. This study was financed by the Coordenação de Aperfeiçoamento de Pessoal de Nível Superior Brasil – CAPES (M.Z.K. grant BEX13621-130) and the Wellcome Trust Investigator awards WT100290MA and Z10661/Z/18/Z to RKG. RKG is part of the Wellcome Trust Centre for Cell Matrix Research funded by award Z03128/Z/16/Z.

## AUTHOR CONTRIBUTIONS

M.Z.K., K.S.H. and A.V.M. performed *in vitro* and *in vivo* experiments; M.Z.K. wrote the original draft of the manuscript, M.R.H., R.K.G., K.N.C. and L.V.S reviewed and edited the manuscript, R.K.G. and K.N.C. conceptual design and project supervision, L.V.S provided resources essential to this study.

## DISCLOSURE

The authors declare that the research was conducted in the absence of any commercial or financial relationships that could be construed as a potential conflict of interest.

## Notes

The authors declare no conflict of interest.

### Competing Interest Statement

The authors have declared no competing interest.

