## supplementary figures for "Large Neutral Amino acid uptake and mTOR activation within CD4+ T cells coordinate Type 2 immunity and host resistance to *Trichuris muris*"

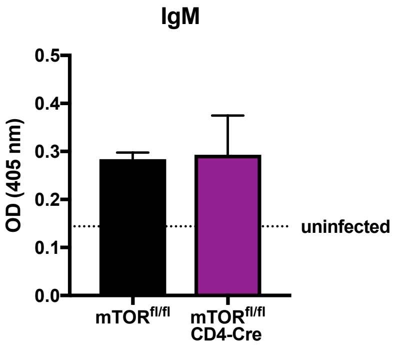


**Fig. S1. Serum levels of parasite-specific IgM in mTOR$^{fl/fl}$ and mTOR$^{fl/fl}$CD4-Cre mice after 30 days of infection.** Mice received 300 infective eggs. *T. muris* E/S was used as antigen. Bar chart shows values for IgM at a 1:20 dilution. Values represent means +/- SEM (n=3).

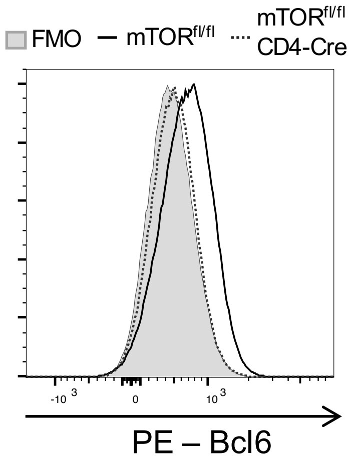


**Fig. S2. Bcl6 in CD4+ T cells MLNs of *T. muris* infected mTOR$^{fl/fl}$ and mTOR$^{fl/fl}$CD4-Cre mice**. MLNs were collected from mice at day 35 post *T. muris* infection (approximately 300 eggs). n=3.

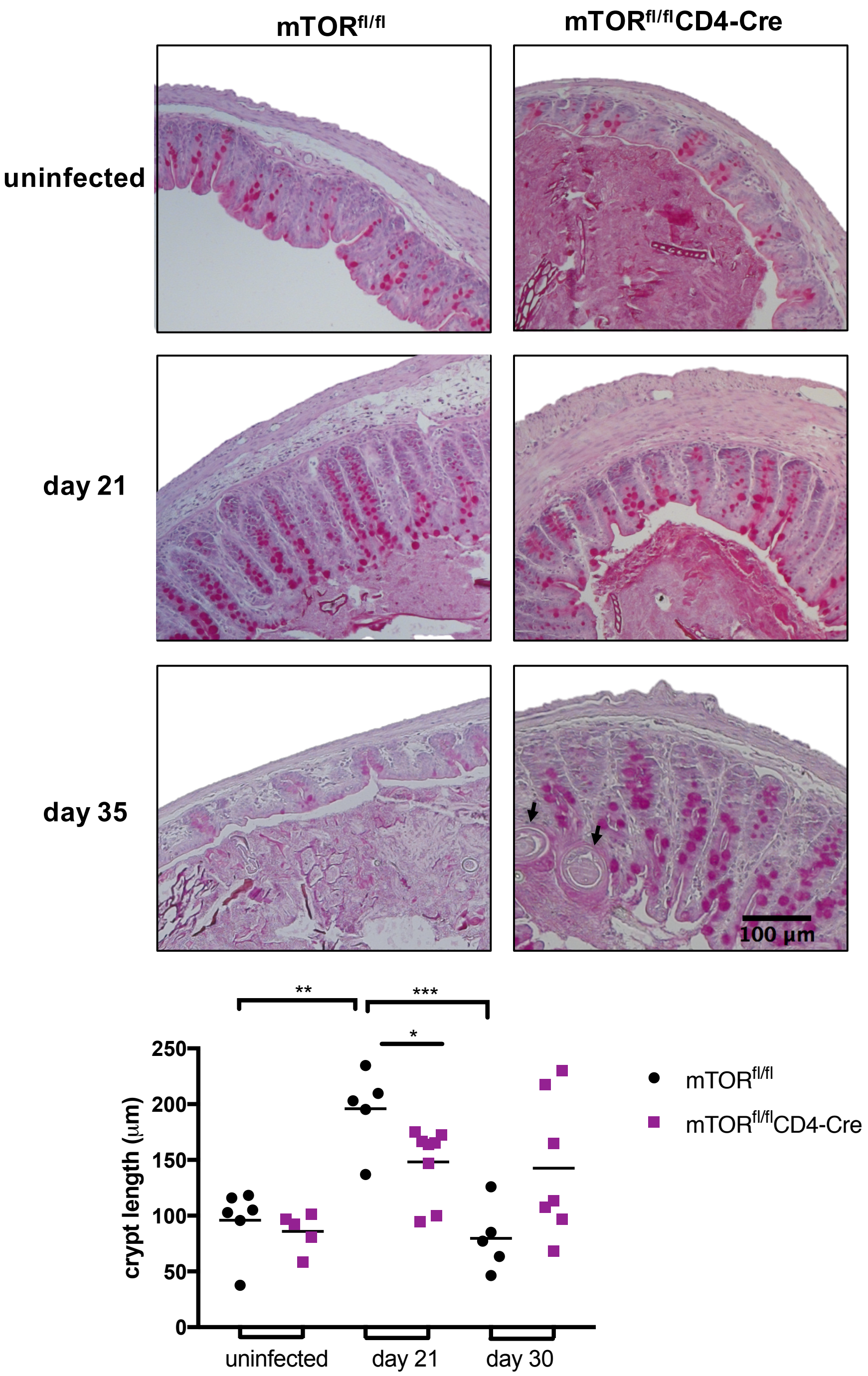


**Fig. S3. Caecal crypt length in naïve and *T. muris* infected mTOR$^{fl/fl}$ and mTOR$^{fl/fl}$CD4-Cre mice**. Mice were culled after 21 or 35 days of receiving a *T. muris* high dose infection. Photographs at x10 magnification were analysed using ImageJ software. Arrows indicate a *T. muris* in cross section. Bars indicate mean (n=5-8). * indicates p value≤ 0.05, **indicates p value≤ 0.01, ***indicates p value≤ 0.001.

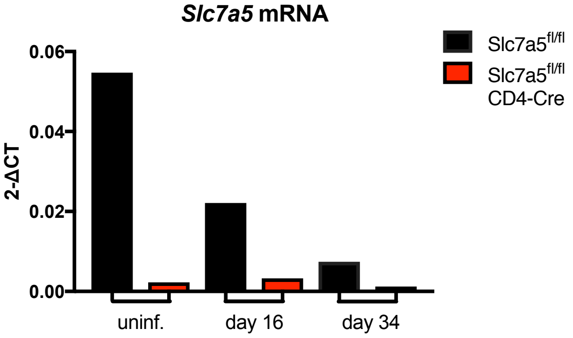


**Fig. S4 Detection of *Slc7a5* in CD4+ T cells from Slc7a5$^{fl/fl}$and Slc7a5$^{fl/fl}$CD4-Cre mice by RT-qPCR.** MLNs were collected after 16 or 34 days of a high dose (300 eggs) *T. muris* infection and from uninfected mice. Cells from 3-5 mice were pooled. Values represent mean CT normalised by b-actin.

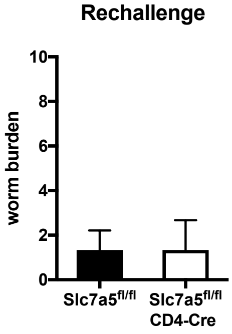


**Fig. S5. Worm burden of Slc7a5$^{fl/fl}$ and Slc7a5$^{fl/fl}$CD4-Cre mice upon *T. muris* high dose re-challenge.** Mice received a high dose of approximately 300 eggs. At day 26 p.i. animals received a second high dose infection. Worm burdens were assessed at day 20 p.i. n=3.

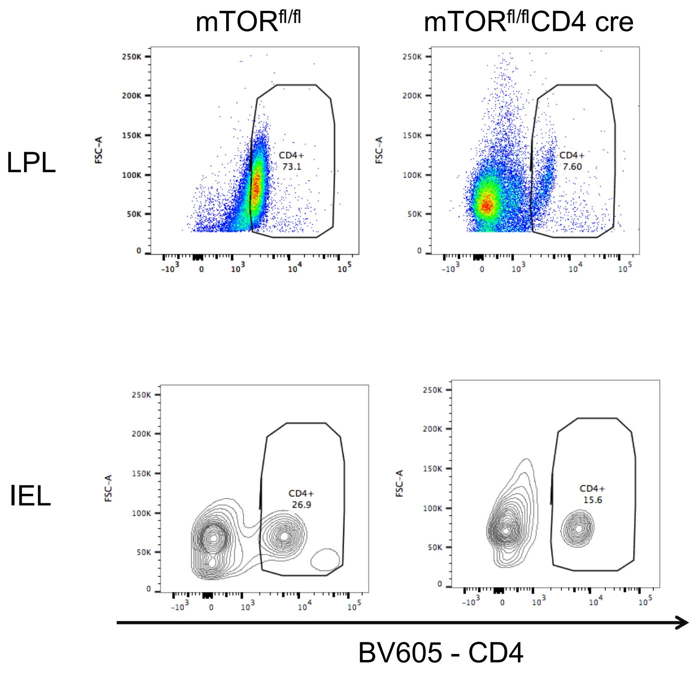


**Fig. S6. CD4+ T cells from the lamina propria and the intraepithelial compartments from mTOR$^{fl/fl}$ and mTOR$^{fl/fl}$CD4-Cre mice infected with *T. muris*.** Cells were isolated from mice at day 21 post *T. muris* infection (approximately 300 eggs). n=5.
